# The pitfalls of negative data bias for the T-cell epitope specificity challenge

**DOI:** 10.1101/2023.04.06.535863

**Authors:** Ceder Dens, Kris Laukens, Wout Bittremieux, Pieter Meysman

## Abstract

Even high-performing machine learning models can have problems when deployed in a real-world setting if the data used to train and test the model contains biases. TCR–epitope binding prediction for novel epitopes is a very important but yet unsolved problem in immunology. In this article, we describe how the technique used to create negative data for the TCR–epitope interaction prediction task can lead to a strong bias and makes that the performance drops to random when tested in a more realistic scenario.

## Main article

Unexpected or unknown biases within machine learning datasets are a common issue that has hindered many well-designed approaches from translating to real-world applications, despite seemingly generalizable performance achieved during model development and evaluation. A well-known example of this issue is a classifier that was trained to identify malignant skin lesions, but ended up relying on the presence of a measuring ruler in the images due to the bias present in the training data^1,2^. However, the presence of data bias is not always obvious. Multiple cases have been reported where specific demographics, such as gender, skin type, ethnicity, or socio-economic status, were underrepresented in the data, leading to unexpected performance differences between different subpopulations and potentially delaying access to care^3,4^. Indeed, as algorithmic approaches become increasingly more advanced and datasets grow larger and are necessarily compiled using less curation, these issues are becoming more and more commonplace. Even small biases within a dataset often suffice for a machine learning model to overfit on bogus data characteristics and drive its predictive behavior. Crucially, if the same bias persists in any held-out test data, this issue will remain undetected.

The same is true for the T-cell epitope prediction challenge, as recently tackled by Gao et al.^5^ T-cells are a critical part of the adaptive immune system, as they recognize intruders from self, induce immune responses, and retain memory. The recognition of foreign intruders is mediated by their T-cell receptor (TCR). When antigen-presenting cells display short peptides (called epitopes) from pathogens or malignant cells, such as cancer cells, on their cell surface, this TCR is able to bind with them in a specific manner, upon which the T-cell will be activated and the immune response will be triggered.

If we would be able to annotate TCR sequences with their targets, this would unlock myriad applications, ranging from vaccine design and cancer treatments to diagnostics. Because of the central role that T-cells play in the immune system, the TCR repertoire of an individual contains valuable information about the past, present, and future immune state. Using high-throughput sequencing technologies, it is possible to map the sequences of the TCRs within a biological sample, for example from blood of an individual with a specific disorder. However, the number of possible TCR sequences is incredibly large, with a conservative estimate in the range of 10^15^ unique sequences^6^. Consequently, the epitope targets of the vast majority of TCRs are unknown. On the other hand, it is known that the specificity of a T-cell is fully driven by its TCR and its static co-receptors^7^. Therefore, the entire recognition event must be encoded within the TCR sequence and is seemingly a straight-forward prediction problem where the right TCR has to be matched with the right target.

Several methods have shown significant potential in extrapolating from a set of TCRs known to bind a specific epitope, to other TCRs targeting the same epitope^8^. However, the number of epitopes with known TCRs is counted in the hundreds, which is just a drop in the ocean of possible TCR targets. Therefore, zero-shot TCR–epitope annotation—i.e. predicting TCR–epitope binding for novel, unseen epitopes—is currently seen as the ‘holy grail’ of immunology^6^. This requires machine learning methods to actually learn the underlying recognition code of the TCRs, which has turned out to be a substantially harder problem.

An important issue that complicates this challenge is the lack of high-quality negative data. While the experimental methods to determine TCR–epitope pairs have a high specificity, they are hindered by a low sensitivity with a high false negative rate^9,10^. For example, the most common approach using tetramers is well-known to only capture a subset of true TCR–epitope interactions^11^. As a result, the number of true negative pairs in TCR–epitope databases is a small fraction of the known positive pairs, and consequently, negative instances are often generated artificially during the development of TCR–epitope prediction models.

There are two approaches commonly used for generating negative data in the context of TCR–epitope annotation (Fig. 1), neither of which are a perfect representation of the real-world scenario. The first is shuffling the known positive pairs, where each TCR is matched with an epitope to create random combinations that differ from those in the positive data. This relies on the principle that a TCR known to be specific for one epitope is unlikely to be specific for another unrelated epitope. However, because of the limited number of epitopes with known TCRs, it is complex to design a held-out negative dataset using this approach. The second strategy, applied by Gao et al.^5^, is using background TCR data. In this case, epitopes from known positive samples are paired with random TCRs from a background set, which is often obtained from a broad sequencing experiment without epitope specificity. However, multiple studies have shown that this second approach confers a substantial bias in the dataset. The study by Moris et al. used a decoy dataset that was generated by replacing each unique epitope with a random amino acid sequence of the same length, removing any chance of true binding^12^. When generating negatives from a background TCR data set, a performance better than random was achieved, demonstrating that the negative data contained a bias that caused a difference between positive and negative CDR3 sequences independent of the paired epitope. Similarly, the study by Grazioli et al. showed that using a background data set to generate negatives for the few-shot or majority learning setting leads to sequence memorization and making predictions only based on the CDR3 sequence, without considering the epitope^13^. The cause of these problems is that the negative pairs and positive pairs are derived from different experiments, performed by different labs, and often even in a different part of the world with different subject ethnicities. Any high-performance machine learning method will be able to capture this bias and utilize it to differentiate between positive and negative samples.

**Figure 1.**
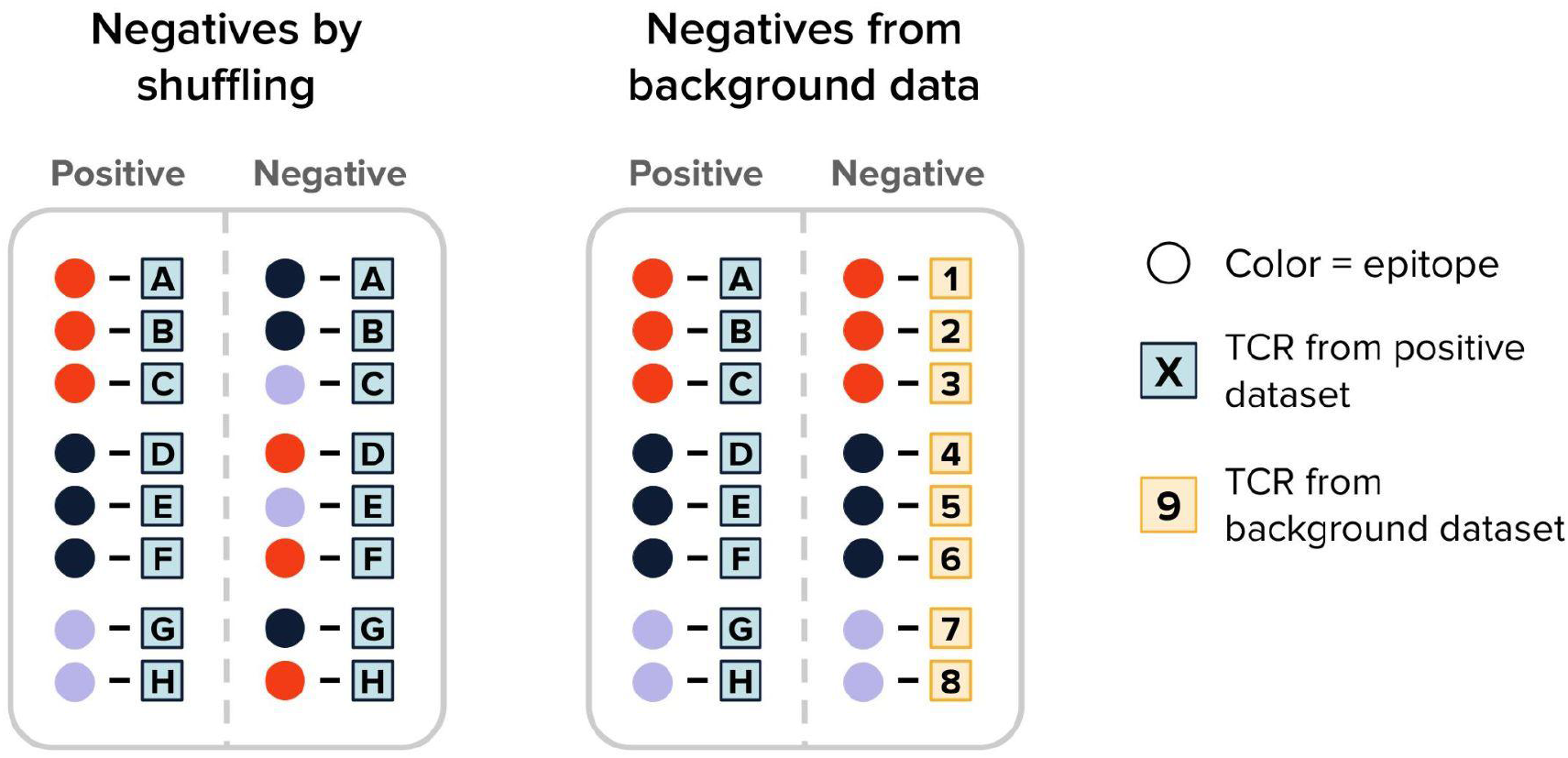
Schematic overview of the two approaches commonly used for generating negative TCR–epitope data. When generating negatives by shuffling (left), the same epitopes and TCR are reused but each TCR is paired with a different epitope. When generating negatives from a background dataset (right), new TCR sequences are paired with the epitopes.

To determine the potential impact of the negative set, we first tested the zero-shot predictions of PanPep using five-fold cross-validation with data generated using the shuffled epitope approach instead of the background TCR approach^12^. PanPep achieved an area under the receiver operating characteristic curve (ROC-AUC) of 54.1% ± 6.4% (mean ± standard deviation) (Fig. 2a), similar to the previously reported ROC-AUC of 54.1% ± 1.9% on this dataset^12^. Note, however, that we did not filter the data to exclude samples or epitope sequences already present in the PanPep training data, with 57.7% of the positive test samples that were part of PanPeps training data and only 3.1% of the test samples that had an epitope not seen during training (see Supplementary Material). As such, although this should have been a relatively easy test, the performance on data with negatives generated by shuffling significantly underperforms the zero-shot ROC-AUC of 70.8% reported originally.

**Figure 2.**
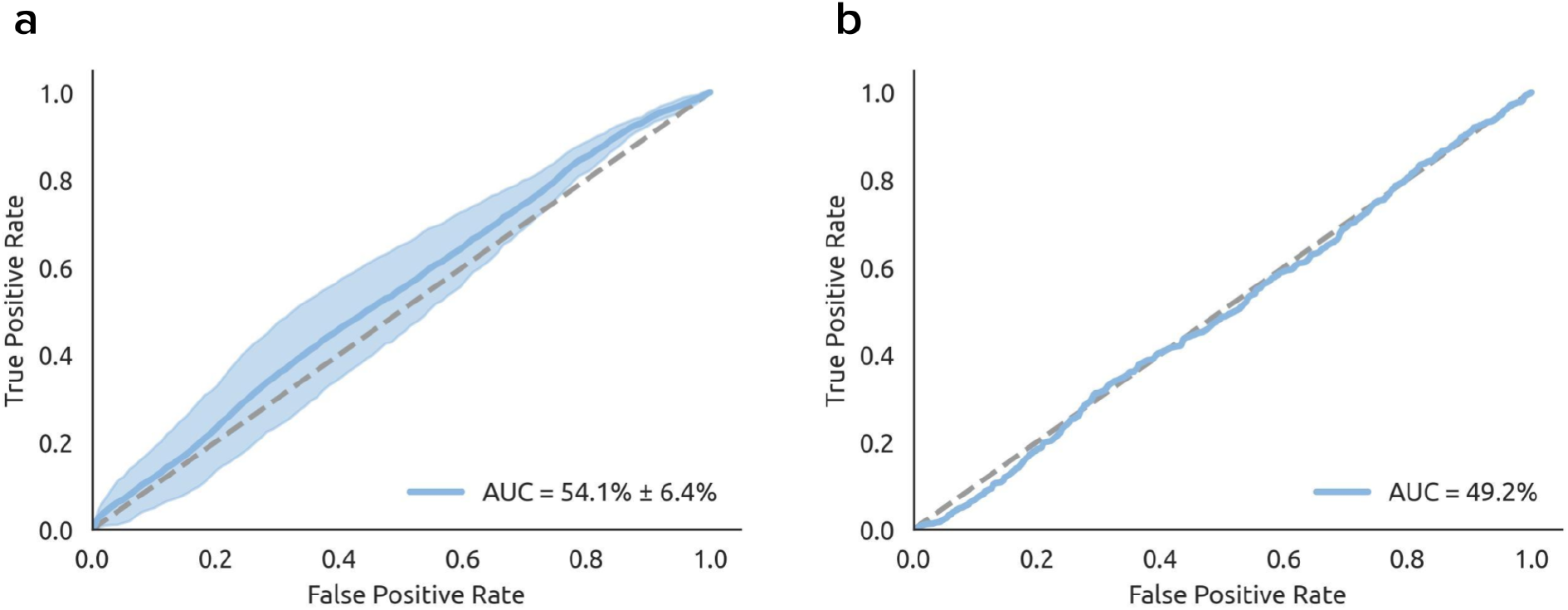
ROC curves of PanPep tested on shuffled negative data. **(a)** Mean ROC curve and standard deviation of PanPep from five-fold cross-validation with data generated through the shuffled epitope approach. The data was not filtered to exclude samples or epitope sequences already present in the PanPep training data. **(b)** ROC curve of PanPep on zero-shot data with negatives generated by shuffling.

Second, we tested PanPep in a true zero-shot setting by using the PanPep zero-shot positive data and generating negative data by shuffling the TCR sequences of these samples. The result is a test dataset that does not contain any samples and epitope sequences already included in the training dataset. On this test dataset, PanPep achieves an ROC-AUC of 49.2% (Fig. 2b), failing to make predictions better than random.

A lack of unbiased labeled data is not unique to the TCR–epitope prediction problem. Similar issues exist within many other fields. For example for speech recognition, a big challenge is the low variability of dialects and accents in the available data^14^. An example with a lack of negative data is anomaly detection, where rare events by definition only occur infrequently^15^. And for protein–ligand binding prediction, a broadly used benchmarking dataset contains a bias in the negative data that makes it easy for machine learning models to distinguish between decoys and binding pairs^16^.

In conclusion, biased data can and will lead to inaccurate and untrustworthy predictions for any machine learning task. This is also the case for TCR–epitope prediction tools trained on biased negative data, as we showed that their performance drops significantly when tested in a more realistic setting. Given the potential advances in healthcare that would arise from accurate TCR–epitope binding prediction tools, we argue that more effort needs to go towards this problem. More data and an unbiased benchmarking dataset are a necessary next step towards prediction models that are reliable in real-world scenarios.

## Competing Interests

KL and PM hold shares in ImmuneWatch BV, an immunoinformatics company.

## Author Contributions

CD performed the study. CD and PM wrote the manuscript. WB, KL, and PM conceived and supervised the study. WB, PM and KL revised the manuscript. All authors read and approved the final manuscript.

## Data and Code Availability

The data and all scripts used to obtain the results are available on Github at https://github.com/PigeonMark/PanPep-Shuffled-Negatives and on Zenodo at https://doi.org/10.5281/zenodo.7798691.

## Materials & Correspondence

Correspondence to Pieter Meysman.

## Notes

https://doi.org/10.5281/zenodo.7798691

### Competing Interest Statement

KL and PM hold shares in ImmuneWatch BV, an immunoinformatics company. No payments or services were received.

