## Supplementary Material: Methods and Data for "The pitfalls of negative data bias for the T-cell epitope specificity challenge"

### Methods

First, PanPep[1] was tested with five-fold cross-validation using the shuffling approach to generate negative samples. PanPep was only tested, not retrained for this task. We used the zero-shot learning prediction setting provided by PanPep.

Second, PanPep was tested on their own zero-shot positive test data, with negative data again generated using the shuffling approach. The zero-shot learning prediction setting provided by PanPep was used.

### Data

The data used for the first test was collected from the August 2019 release of VDJdb[2]. This dataset was filtered on human TCR beta sequences, all spurious CDR3 sequences as defined by VDJdb were removed, only MHC class I entries were kept, all entries originating from the 10x Genomics demonstration study[3] were omitted, the length of the CDR3 and epitope sequences was filtered to lie between 10–20 and 8–11 amino acids respectively, and finally any duplicate sequence pairs were removed. The data was downsampled to have at most 400 samples per epitope and negatives were generated by shuffling[4].

From this data, five zero-shot train/test cross-validation splits were generated but only the test splits were used to test PanPep. Because both methods use partly the same sources for their positive samples, we calculated the overlap between the positive samples in the test sets we generated and the PanPep training data. All numbers are an average over the 5 cross-validation folds. 773.6 out of 1340.4 (57.7%) of all positive test samples were already in the PanPep train dataset. 1298.8 out of 1340.4 (96.9%) of all positive test samples have an epitope that was already in the PanPep train dataset. 16.2 out of 23.6 (68.6%) unique epitopes in the test dataset were already in the PanPep train dataset.

For the second test, the zero-shot positive test dataset from PanPep was used. As described previously, this data was collected from IEDB[5], VDJdb[2], PIRD[6], and McPas-TCR[7]. The data was filtered on human TCR beta sequences, only HLA-I entries were retained, the records in VDJdb with a confidence score of 0 were removed and only the high-confidence binding TCRs in the PRID dataset were retained. The train dataset was created by only using the samples with a peptide with at least five binding TCR records. The remaining samples were used for the zero-shot test dataset. We added negatives by shuffling the TCR sequences. The resulting test dataset is truly zero-shot, none of the samples were already in the PanPep train data and none of the samples had an epitope also present in the PanPep train data.
